# An active traveling wave of Eda/NF-kB signaling controls the timing and hexagonal pattern of skin appendages in zebrafish

**DOI:** 10.1101/2023.04.10.536269

**Authors:** Maya N. Evanitsky, Stefano Di Talia

## Abstract

Periodic patterns make up a variety of tissues, including skin appendages such as feathers and scales. Skin appendages serve important and diverse functions across vertebrates, yet the mechanisms that regulate their patterning are not fully understood. Here, we have used live imaging to investigate dynamic signals regulating the ontogeny of zebrafish scales. Scales are bony skin appendages which develop sequentially along the anterior-posterior and dorsal-ventral axes to cover the fish in a hexagonal array. We have found that scale development requires cell-cell communication and is coordinated through an active wave mechanism. Using a live transcriptional reporter, we show that a wave of Eda/NF-κB activity precedes scale initiation and is required for scale formation. Experiments decoupling the propagation of the wave from dermal placode formation and osteoblast differentiation demonstrate that the Eda/NF-kB activity wavefront times the sequential patterning of scales. Moreover, this decoupling resulted in defects in scale size and significant deviations in the hexagonal patterning of scales. Thus, our results demonstrate that a biochemical traveling wave coordinates scale initiation and proper hexagonal patterning across the fish body.

## Introduction

Skin appendages are a key feature of vertebrate anatomy, and have a myriad of functions and forms, from hair in mammals to feathers in birds^1–3^. Formation of these structures is complex and includes periodically and dynamically delivered patterning sequences. The emergence of periodic patterns is a fundamental process in development and drives the formation of a variety of tissues, both internal and external. Periodic patterns can arise from a homogeneous field of cells through symmetry breaking mechanisms such as self-organization or changes in gene expression at the cellular level. Experimental access to this dynamic sequence is a challenge that often is limited by a lack of direct access to the process^2,3^. Many of the mechanisms that regulate periodic patterning occur early in development, when tissues such as the skin or gut are inaccessible. Mammals and birds, two common model systems to study skin development, develop *in utero* or *in ovo*, making longitudinal studies difficult^4^. Here, we develop a system for exploration via live imaging of zebrafish scales as they develop.

Zebrafish scales are skin appendages that cover the fish in a highly organized, hexagonal array. Their development process is similar to other skin appendages and the pathways that regulate this are conserved across vertebrates. Unlike skin appendages in other vertebrate systems, zebrafish scales develop relatively late in ontogeny, approximately four weeks post-fertilization, and are transparent, providing a unique opportunity for longitudinal studies examining tissue growth and patterning^1,5^.

Zebrafish scales consist of osteoblasts surrounding a mineralized bone matrix and mirror mammalian dermal bone development by forming via intramembranous ossification^6,7^. Scales do not appear all at once over the fish, but sequentially in a spreading wave. The first scales begin forming in two locations, one in the trunk of the fish and one more anterior on the belly, then spread out along the anterior-posterior and dorsal-ventral axes until the entire fish is covered (Figure 1a-d). Recent studies have begun to identify the molecular pathways involved, but exactly how this process is regulated remains unclear^1,5,8^.

While bony scale development is unique to fish, the pathways that coordinate this process, namely ectodysplasin A (Eda), Wnt, and Fgf, regulate skin appendage formation across vertebrates^4,9^. Eda is a member of the TNF family and binds to activate its receptor, Edar. This leads to NF-κB then translocating into the nucleus to activate target genes, including Wnt, Shh, and Fgf^10^. In zebrafish, *Eda* is expressed in dermal fibroblasts and null mutations in *eda* lead to a complete loss of scales. On the contrary, *edar* is expressed in epidermal cells above the scale papillae^1,11^.

The process by which cells migrate and differentiate to form skin appendages is likewise conserved. In zebrafish, mesenchymal cells in the dermis condense to form placodes. At the same time, signals from the epidermis trigger growth and differentiation in these cells. The placode then develops into a layer of osteoblasts surrounding a mineralized matrix of bone^5,8,12^ (Figure 1g,h).

**Figure 1:**
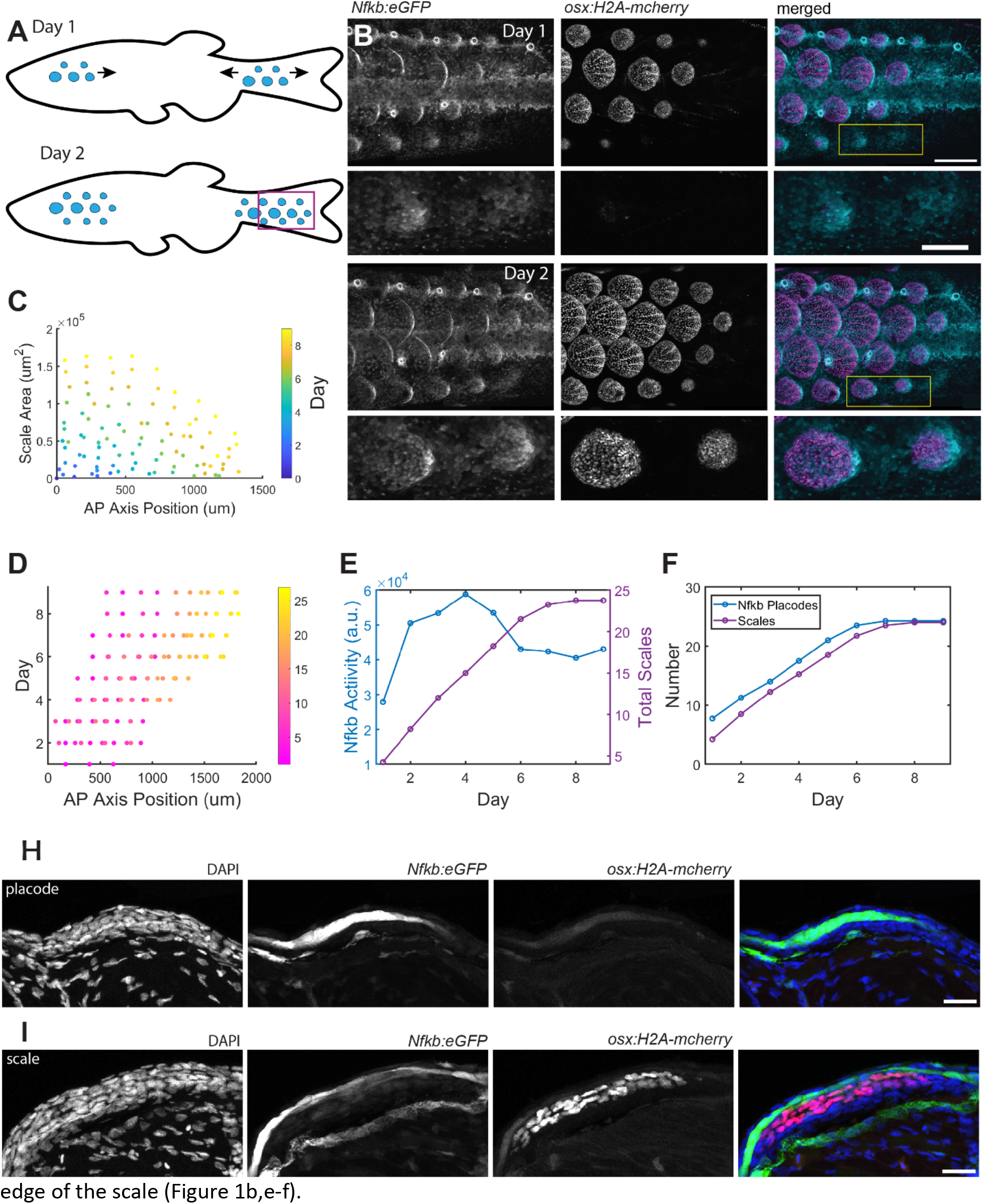
Scale development and NF-κB activity proceed in a wave-like pattern. **A** Model of scale development. Scale formation begins along the midline and spreads in a wave across the body. Rectangle denotes area imaged in panel **B** and subsequent experiments. **B** NF-κB activity (left, cyan) precedes scale formation (marked by osx:H2A-mCherry). *NfκB:eGFP* signal can be seen in a horizontal stripe along the fish midline, lateral line cells, and scale placodes, which differentiate into osteoblasts (middle, magenta) at day 2, scale bar: 250μm. Yellow rectangle denotes insets in lower panels, scale bar: 100μm **C** Quantification of scale size versus location along the anterior-posterior axis. Zero marks start of images taken in the trunk region (see rectangle in panel **A**). New scales start small, then increase in size as new scales are added to the posterior, n=4. **D** Quantification of day of scale development versus scale location showing that new scales form posterior to existing scales, n=4. Color bar refers to scale identification number used to track individual scales. **E** NF-κB activity and number of scales during scale development. NF-κB activity starts low when few scales are present, then increases as more scales and placodes form before leveling off as scale development finishes, n=4. **F** Number of NF-κB placodes and scales during scale development, n=4. **G** Representative image of cross-section of a nascent NF-κB placode, prior to osteoblast differentiation (marked by lack of *osterix* expression) scale bar: 25μm. **H** Representative image of cross-section of a nascent scale with differentiated osteoblasts, scale bar: 25μm. NF-κB is active in the epidermis over the placode and scale; the GFP signal below the scale is background from the midline (see panel **B**).

Although much has been discovered regarding the formation of individual hair follicles or feathers, establishing the regulators of the temporal and spatial patterning has remained elusive. Signaling waves are emerging as a general mechanism by which developing tissues are organized. In systems that work in a periodic and sequential manner, waves contribute to directionality, sequentiality and spacing of the patterns^9,13^.

During skin development, a tissue-wide priming wave could shift a homogenous system of cells into a state to allow for symmetry breaking and facilitate the spatial and temporal organization of the pattern. This priming wave could be either an active or passive wave. Active waves depend on diffusion and biochemical feedback to travel across a tissue while passive waves are kinematic and are controlled by pre-determined spatial delays (for example, delays imposed by nonuniform oscillators)^14,15^. While early studies have suggested that the wave which patterns feather placodes might be passive, more recent work has shown that the morphogenetic signals traveling ahead of placode formation are in fact active waves^9,16,17^. As these early experiments were done in chicken skin explants without the benefit of live molecular markers, it is likely that the signals regulating feather formation were not interrupted. Due to the dynamic nature of these waves, the timing of experiments to interrupt wave propagation is crucial. Longitudinal studies with molecular markers are also necessary for distinguishing passive and active waves^3,16^.

One of the key regulators of skin appendage development, Eda, is expressed as a wave in the epidermis and drives feather patterning^9^. Eda upregulates Fgf20 expression, which acts as a chemoattractant to promote dermal cell condensation. These condensates in turn activate more Fgf20 signaling through mechanical feedback. Dermal cell aggregation is driven by cell migration and occurs in a wave-like pattern. Although Eda and Fgf20 promote the formation of cell condensates in the dermis, it remains unclear whether the formation of condensates precedes the wave of Eda expression or vice versa and what the dependencies among these signals are^9^. Moreover, it has also been suggested that the formation of feather placodes is initiated by mechanical signals rather than biochemical ones^18^. Due to the lack of live imaging techniques and longitudinal studies, it has been difficult to dissect which processes precede and drive one another and the precise relationship between these processes remains unclear. The ability to study scale formation longitudinally in living animals using transgenic reporters and pharmacological perturbations provides a unique opportunity to dissect these dependencies in the formation of zebrafish scales.

## Results

The transcription factor NF-κB acts downstream of Eda signaling and is important for skin appendage development^1,10,19^. However, the dynamics of NF-κB activity in appendage development have yet to be quantitatively described. To that end, we used an osteoblast-marker to label scales (*osx:H2A-mcherry*)^6^ and a previously established transcriptional reporter transgenic line (*Tg(6xHsa.NFKB:EGFP)*^20^, hereafter *Nfkb:eGFP*, to visualize NF-κB activity and its dynamics during scale development. This line features six NF-κB binding sites fused to a minimal promoter to drive GFP expression when NF-κB is in the nucleus. We focus on the mechanisms of scale formation near the caudal fin (Figure 1A). By live imaging skin patterning every 24 hours, we found that NF-κB is transcriptionally active in the epidermis prior to osteoblast differentiation. While NF-κB acts downstream of multiple signaling pathways and labels multiple tissues, including the fish lateral line cells and immune cells in the skin, NF-κB activity can be specifically detected in a primordium that precedes the specification of scale osteoblasts by about a day. As scales grow and mature, NF-κB activity becomes restricted to the epidermis overlying the posterior

Using this line as a reporter, we investigated whether scale development and Eda/NF-κB activity traveled as an active or passive wave. In the case of an active wave, communication between cells is required to spread the signal throughout a tissue, while in a passive wave, each scale is specified autonomously in a predetermined temporal pattern. These two models can be differentiated by their behavior when a barrier is introduced to the system^14,21^. As active waves require a (diffusible) signal for cell-cell communication and some form of positive feedback to propagate across a tissue, the signal will be unable to pass through the barrier and scale development will be halted. A passive wave, on the other hand, does not require cell-cell communication as it is controlled by pre-determined delays and thus the wave can cross the barrier and scale development will continue unaltered (Figure 2a).

**Figure 2:**
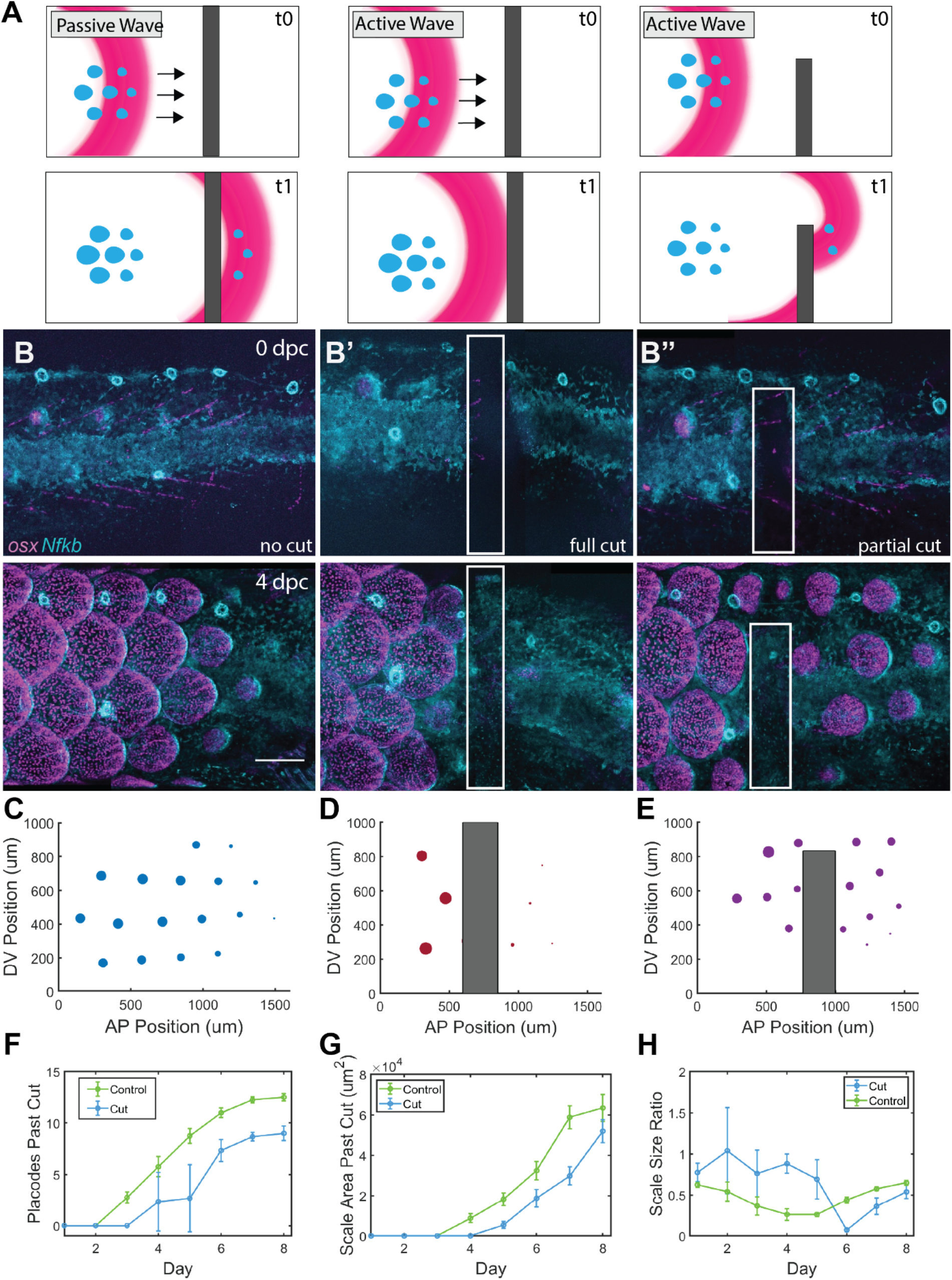
Scale development results from an active wave mechanism. **A** Schematic of phase wave (left) and active wave (middle, right) propagation in response to a barrier blocking cell-cell communication. Blue objects represent scales and pink crescent is the traveling wave. Top row: time point 0, bottom row: time point 1. **B** Representative images of wild-type fish during scale development, cyan: *Nfkb:eGFP*, magenta: *osx-H2A-mcherry*. Dpc: days post-cut. **B’,B’’** Small incisions were made in the skin of the fish to block cell-cell communication. The white box outlines the area of the cut, scale bar: 250μm. **C,D,E** Quantification of scale location and size in uninjured fish, full cut, and partial cut; size of dots is proportional to scale area. **F** Cuts block the formation of placodes posterior to the cut site. Placodes were quantified posterior to cut site (new placodes formed after the cut healed at day 4-5) and in a comparable region in uninjured controls (pchi2=7.51E-21). **G** After cuts healed, growth of individual scales is comparable between injured fish and controls (pchi2=2.18E-10). **H** Size ratio between neighboring scales at the posterior edge of the wave was calculated, smaller scale ratio indicates a greater difference in scale size between neighboring scales (pchi2=4.99E-21) n=7.

Previous studies found that cutting explanted chick embryo skin just ahead of newly formed placodes did not impede the formation of new placodes, suggesting that in that system the wave might be passive^16,22^. However, the timing of these experiments is key, and these studies predated the existence of live molecular markers for feather formation, so it is possible that the cut might have been introduced after the initiating wave had already traveled across the cut region. Consistent with this scenario, subsequent experiments introducing incisions further from the initiating line of feather placodes blocked the formation of new placodes, indicating the wave might in fact be an active one^17^.

To determine whether NF-kB waves during scale formation in zebrafish are active or passive, we made a shallow incision with a razor blade along the dorsal-ventral axis of juvenile fish to interrupt cell-cell communication. Cuts were made posterior to *Nfkb:eGFP* activity and scale formation at a stage in which scale induction proceeds anterior to posterior in the trunk of the fish (Figure 2b). Fish were imaged once a day until the cut fully healed.

In each fish, cuts caused a significant delay in the formation of new scales. Prior to cut healing, no scales formed posterior to cuts that bisected the entire anterior-posterior axis. Although the injury did impact the growth of scales in the cut site, scales anterior the cut continued to develop and grow normally. Only after the cuts were fully healed did new scales begin to form posterior to the cut site. We also performed cuts that did not completely bisect the anterior-posterior axis, allowing signaling to propagate above the cut. Unlike the full cuts, NF-κB activity and scale development proceeded around the cut and continued in the expected pattern (Figure 2b-h). Importantly, cuts made parallel to the anterior-posterior axis did not impact scale development outside of the cut site (Supplementary Figure S1), indicating that the delay in scale formation is not triggered by a global injury response.

These results suggest that new scales only form in response to a signal from neighboring cells, rather than being instructed by a pre-existing pattern. If the cuts had no effect on scale development outside of the injury site, this would suggest that a cell-autonomous mechanism dictated the timing of scale development. The fact that the experimental cuts impeded the traveling of NF-κB activity and subsequent scale development suggests scales develop via an active wave mechanism. Furthermore, as scales only formed following the emergence of NF-κB activity, this suggests that NF-κB may itself act as the primary factor instructing scale initiation.

To further examine the role of NF-κB in this system, we used the pharmacological inhibitor Bay11-7085 to manipulate NF-κB activity during scale development^19,23^. Fish expressing *Nfkb:eGFP* and *osx:H2A-mcherry* were imaged and treated with Bay11-7085 over the course of several days. We found that Bay11-7085 caused a reduction in *Nfkb:eGFP* signal and a significant reduction in the rate of scale formation and growth (Figure 3b-d). A transient treatment with Bay11-7085 revealed that scale development stopped during treatment but recovered following drug washout and the return of NF-κB activity. Moreover, this temporary inhibition did not significantly affect the patterning of scale development as new scales formed sequentially in the pre-established pattern set prior to the inhibition of NF-κB (Figure 3e-g).

**Figure 3:**
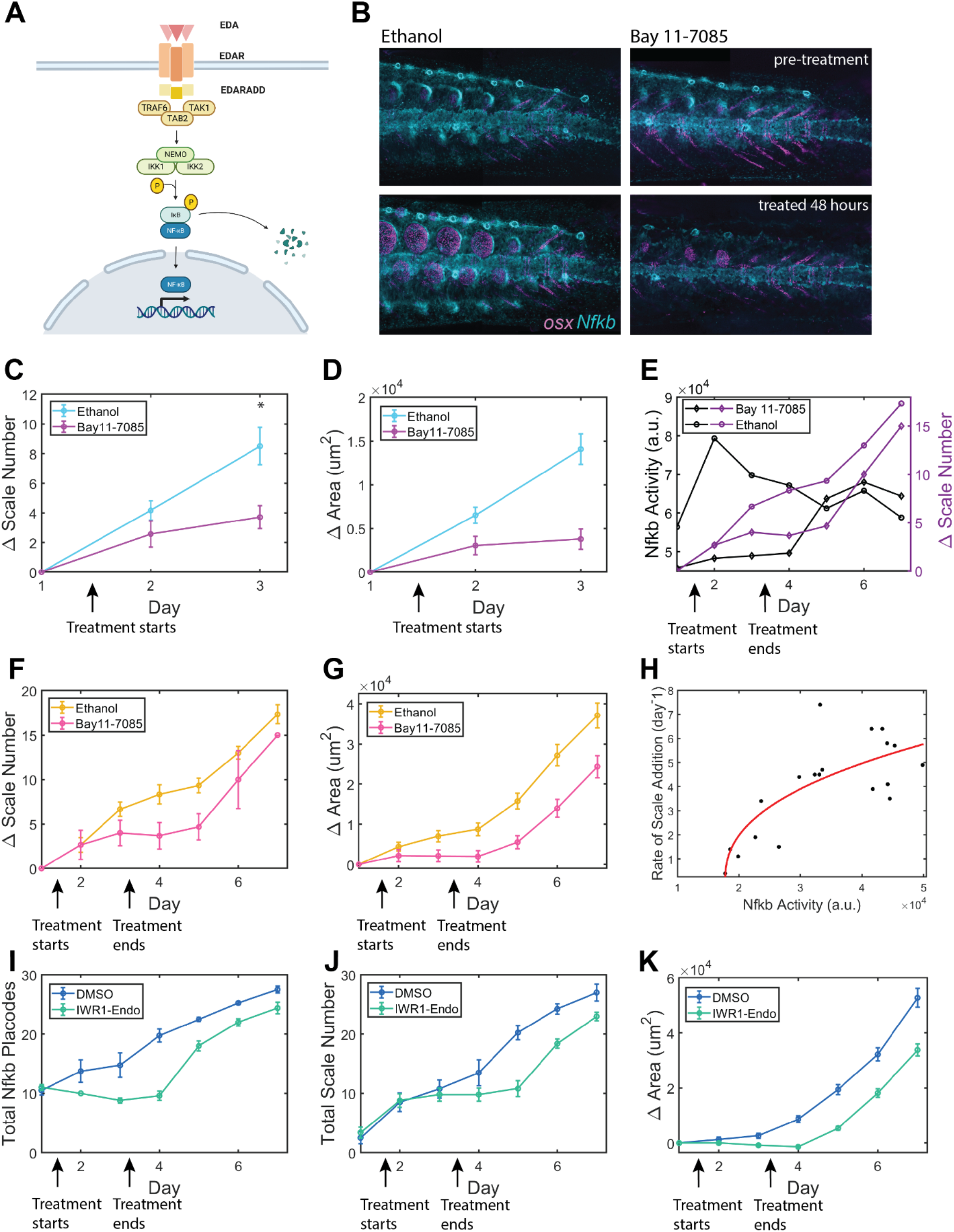
NF-κB activity regulates scale development. **A** Diagram of Eda signaling pathway. Eda binds to its receptor, EDAR, which forms a complex to activate the IKK complex. IKK targets IkB for degradation, releasing NF-κB to enter the nucleus. **B** Representative images of *Nfkb:eGFP* (cyan);*osx:H2a-mcherry* (magenta) fish before and during treatment with NF-κB inhibitor, Bay 11-7085. Treatment causes a loss of *Nfkb:eGFP* signal. **C,D** NF-κB inhibition slows the formation of new scales (day 3 T-test p=0.0038) and the growth of individual scales (day 3 T-test p=1.4964e-05), n=13, error bar = SEM. **E** NF-κB activity (black lines, left axis) correlates with scale formation (purple lines, right axis). During treatment, NF-κB activity and scale formation are reduced. After washout, NF-κB returns at day 5 and scale addition increases at day 6. **F,G** Fish were treated with Bay 11-7085 from day 1.5 (post-imaging) until day 3.5. During treatment, addition of scales (pchi2 = 0.001) and scale growth (pchi2 = 1.58E-10) was reduced and resumed as normal after washout, n=6. **H** Rate of scale addition correlates with NF-κB activity. Rate of scale addition per day was calculated using a linear regression fit, n=19. **I, J,K** Fish were treated with Wnt inhibitor, IWR1-Endo, from day 1.5 to day 3.5. During treatment, the total number of Nfkb placodes (pchi2 = 2.31E-24) and scales (pchi2 = 1.3476e-11) were reduced or did not increase. Scale growth was also inhibited (pchi2 = 2.94E-39).

In an active model in which NF-kB contributes to the mechanism triggering the wave, the speed of the wave depends on the time scale of self-sustained NF-κB activation. In a passive wave model, however, NF-κB does not participate in wave propagation, as the wave is predetermined by an upstream factor, and thus the speed of the traveling wave would be independent of NF-κB activation kinetics. We found a significant correlation between NF-κB activity and the rate of scale formation (used here as a proxy for wave speed), arguing NF-κB activity contributes to the propagation of the wave. This correlation and the observation that NF-κB activity precedes new scale formation suggest that NF-κB activity plays a central role in the timing of scale development (Figure 3h).

As NF-κB activity propagates as an active traveling wave to induce scale formation, we suggest that NF-κB promotes downstream signals promoting both the activation and inhibition of the Eda/NF-κB pathway to create a transient signal that can move across the skin^24,25^. As such, active NF-κB in a cell must be able to promote the pathway in a neighboring cell, likely through the production and release of extracellular ligand(s). Since the developmental ligand Eda is upstream of NF-κB (Figure 3a), it is natural to consider the possibility that it is involved in the positive feedback driving the observed NF-κB activity wave. Consistent with this, a wave of Eda expression has been observed during feather formation. However, how this wave is propagated is unclear^1,9^. Since in zebrafish, NF-κB is active in the epidermis, where the Eda receptor is expressed (Edar), while Eda is solely expressed in the dermis^1,11^, it is unlikely that NF-κB can directly regulate Eda transcription. Thus, a simple feedback loop of Eda activating NF-κB to produce more Eda is improbable. This suggests that the wave is propagated through an intermediary molecule that is expressed in the epidermis and active in the dermis. A strong candidate to participate in the wave is Wnt/β-catenin signaling, which is required for osteoblast differentiation, is active in the epidermis and dermis during scale development, and is capable of activating Eda in other systems^1,26,27^.

To investigate the potential role of Wnt/β-catenin signaling in this process, we used the pharmacological inhibitor IWR-1-endo, which has been shown previously to affect scale development in juvenile zebrafish^1,26,28^. Similar to blocking NF-κB activity, inhibition of Wnt signaling halted scale formation and the formation of new NF-κB placodes. Upon release of inhibition, scale and placode development proceeded as normal (Figure 3i-k), as observed following NF-kB transient inhibition. The similar effects of inhibition of Wnt and Eda/NF-kB signaling on scale formation suggests that they might cooperate in the propagation of a signaling wave that triggers a wave of scale patterning.

In addition to Wnt and Eda, Fgf signaling is also required for scale development^1,11,29^. A null mutation in Fgfr1 results in abnormally large scales, but in combination with an *fgf20* mutation, scales are smaller and fewer, suggesting that multiple Fgfs and Fgfrs regulate scale development^29^. In other systems Fgf20 has been shown to act downstream of Eda and to promote mesenchymal aggregation and placode formation^4,9,10^. Fgf20 may act as a chemoattractant to drive dermal cell condensation, which occurs in a wave-like pattern^8^. Using a transcriptional reporter of *fgf20a* expression^6^, we found that *fgf20a* expression closely mirrors NF-κB activity. *Fgf20a* is on early in the scale placode prior to *osterix* expression and then becomes restricted to the exterior edge of the scale as the tissue grows. Following inhibition of NF-κB, formation of new *fgf20a* positive placodes stopped, indicating NF-κB is required to activate Fgf20a expression (Figure 4a-c).

**Figure 4.**
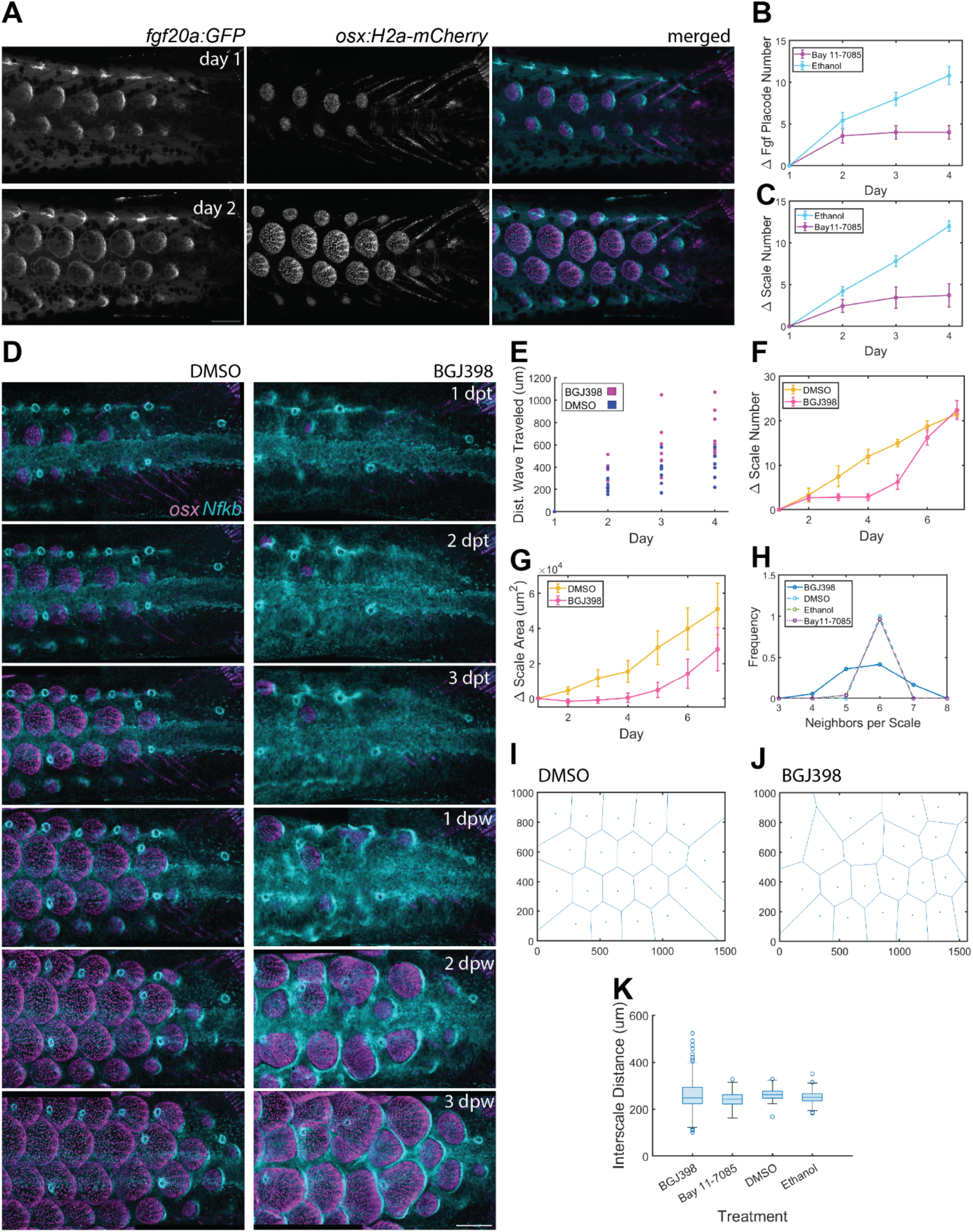
A sequential wave of NF-κB activity times scale formation and is required for hexagonal patterning. **A** Wild-type fish expressing *fgf20a:GFP* and *osx:H2a-mcherry* during scale development. Fgf20a expression precedes scale formation. **B,C** Fish expressing *fgf20a:GFP* and *osx:H2a-mcherry* were treated with NF-κB inhibitor, Bay 11-7085 starting at day 1.5. Number of Fgf20a placodes (pchi2 = 1.40E-08) and scales (pchi2=3.2232e-09) formed were quantified, n=12. **D** Representative images of fish expressing *Nfkb:GFP* (cyan) and *osx:H2a-mcherry* (magenta). Fish were treated with pan-FGFR inhibitor, BGJ398 until 3dpt; dpt: days post-treatment start, dpw: days post-washout, scale bar: 250 μm. **E** Quantification of distance NF-κB wave traveled during treatment with BGJ398, pchi2 > 0.05, n = 16. **F, G** Quantification of change in scale number (pchi2= 7.67E-12) and scale size (pchi2= 1.64E-04) during BGJ398 washout. Fish were treated starting at day 1.5 until day 3.5, n=9. **H** Quantification of scales directly adjacent to a single scale at day 7 of washout experiments for BGJ398 and Bay11-7085 experiments. **I,J** A Voronoi tessellation was constructed using scale centroids from day 7 of BGJ398 washout experiment. Boundary Voronoi cells were not used for quantification. **K** Quantification of distance between scales at day 7 of washout experiments for BGJ398 and Bay11-7085 experiments.

Next, we used the pan-Fgfr inhibitor BGJ398^1,21^ to analyze the contributions of Fgfr signaling to both dermal condensation and wave propagation. Treatment with this inhibitor completely blocked scale development as anticipated, but, similar to observation in feather formation^9^, did not block the Eda/NF-κB activity wave from traveling. The Eda/NF-κB activity wave not only continued to travel along the anterior-posterior axis but exhibited an ectopic pattern. Instead of distinct placodes of NF-κB transcription, we observed a continuous front of reporter expression (Supplementary Figure S2). This suggests that Fgf signaling is not required for the propagation of the Eda/NF-κB wave and may play a role in activating the signals responsible for turning off Eda and NF-κB (Figure 4d,e).

Since the Eda/NF-kB activity wave continues to travel in the absence of Fgf signaling but no obvious placodes could be detected, we used a transient treatment with the BGJ398 to test the effects on patterning of decoupling wave propagation and placode formation. Following drug washout, the extended propagation of the Eda/NF-κB activity wave led to significant changes in scale patterning. Scales formed post-washout arose more synchronously, resulting in a similar number of scales compared to controls (Figure 4f,g). Additionally, the spatial pattern of these new scales was significantly altered as well (Figure 4h-k). Rather than a strict hexagonal arrangement, there was greater variation in the number of neighbors each scale had and the spacing between scales. This result suggests that the sequential timing of scale development is required to properly pattern scales. Thus, coupling the traveling Eda/NF-κB activity wave to dermal condensation and scale formation is necessary to facilitate accurate patterning and growth.

## Discussion

Zebrafish scales are an ideal system to study skin patterning as their external development and the use of transgenic reporters and live imaging are conducive to dynamical analyses. Developing scales form relatively late in ontogeny. Thus, they can be easily imaged to conduct long term experiments and manipulating the processes that drive scale development is unlikely to affect the formation of other critical structures, which might in principle confound the analysis of these perturbations. As the pathways and developmental mechanisms that regulate skin appendage development are conserved across vertebrates, this system can improve our quantitative understanding of how these pathways lead to patterning.

Here, we have shown that skin appendage development in zebrafish is coordinated by the propagation of a biochemical wave. We have found that an active wave of NF-κB activity regulates the spatial and temporal patterning of scale development. Interfering with cell-cell communication blocked the propagation of Eda/NF-κB activity and scale development. Both NF-κB and Wnt activity are required for scale development and propagation of the wave. Fgf signaling, while required for scale development, is not involved in how this wave travels. Although it is possible to drive scale development through ectopic Fgf signaling in the absence of Eda, this results in aberrant scale patterning and ectopic osteoblast formation^1^. Since Fgf signaling is required for the formation of the dermal placodes that precede scale formation, these results demonstrate that the signaling waves and the cellular processes that lead to placode formation (for example, migration) must be tightly coordinated to properly pattern scale development.

Our results suggest a mechanism by which the wave of Eda travels across the skin to drive this process. As NF-κB is active in the epidermis and Eda is expressed in the dermis, it is unlikely that NF-κB directly activates transcription of Eda^1^. Thus, NF-κB likely induces Wnt, which can then diffuse to the dermis to trigger expression of Eda. Eda can in turn diffuse to the epidermis to activate NF-κB and complete the positive feedback loop. At the same time, NF-κB also induces *fgf20a* expression in the epidermis, which can signal to drive dermal cell aggregation, which possibly promotes the inhibitors responsible for turning off Eda/NF-κB signaling late in scale development. While these inhibitors have yet to be identified, Bmp signaling has been shown to inhibit Eda and could be acting to block scale formation^4,9,10^.

In recent years, signaling waves have emerged as a common mechanism for the regulation of developmental processes^21,25,30^. Waves can facilitate the rapid propagation of biochemical signals and help organize cellular responses across large tissues. In cellular systems, this process involves ligand release and often *de novo* production mediated by transcription. The timescales of such processes dictate the speed of the wave. In systems in which wave propagation involves transcription, the typical speed of traveling waves is about 10 microns/hour, that is the wave travels from one cell to the other in about an hour^14^. Consistent with this idea, we found that the wave of Eda/NF-κB travels about 200 microns in a day, suggesting that the wave is likely driven by the transcription and release of ligands at the wavefront.

In summary, we have shown the efficacy of using zebrafish as a model for vertebrate skin appendage development and demonstrated that a wave of Eda/NF-κB activity is required to form the precise hexagonal pattern in scale development. We decoupled wave propagation from the formation of the placodes by transiently inhibiting Fgf signaling. Consequently, the Eda/NF-κB wavefront travels across a large region of tissue but no placode forms. Following washout of the inhibitor, a large region of the tissue becomes competent to form placodes simultaneously. As a result, several scales form at the same time but they do in a less precise and regular pattern. Thus, we propose that a biochemical wave is upstream in the patterning process and that its coupling with the mechanochemical process of placode formation ensures that sequential formation of scales proceeds in precisely controlled hexagonal array.

## Methods

### Fish Husbandry

Zebrafish of Ekkwill and Ekkwill/AB strains were maintained between 26-28.5°C with a 14:10 hour light:dark cycle. Juvenile fish aged between 3.5-5 weeks post-fertilization were used for experiments. Both males and females were used for experiments and randomly assigned to control and experimental groups. As scale development does not start at an exact age, all fish were checked for the onset of placode or scale development before use in experiments. For each experiment, control and experimental groups were taken from the same clutch. All fish experiments were approved by the Institutional Animal Care and Use Committee at Duke University and followed all the relevant guidelines and regulations. Transgenic lines used in this study were Tg(osx:H2A-mCherry)_pd310_^6^, Tg(fgf20a_EGFP_)/HGN21A^31^, Tg(6xHsa.NFKB:EGFP)^20^.

### Live Imaging

Live juvenile fish were imaged using a Leica Sp8 confocal microscope and LAS X 2.01.14392 software with a 10X air objective at 1.25X zoom. Fish were anesthetized with 0.015% tricaine (Sigma E10521-50G) in system water and transferred to a 1% agarose bed in a 40mm petri dish. After imaging, fish were revived and placed back in their original tanks. For longitudinal time courses, fish were imaged once every 24 hours for 3-10 days. All images focused on the trunk of the zebrafish, just anterior to the fin. As the area of scale development is larger than the field-of-view of our microscopy setup, multiple overlapping z-stacks (1-3 with variable number of plans) were used. Images were acquired at 1024 × 1024 resolution (0.909-μm pixel size) and a z-step of 0.909 μm. Fluorescent proteins were imaged using the following lasers: *Nfkb:eGFP*, 488nm, *osx:H2A-mCherry*, 561nm, *fgf20a:GFP*, 488nm.

### Skin Injuries

Fish expressing *Nfkb:eGFP* and *osx:H2a-mcherry* were anesthetized using 0.015% tricaine in system water. Once fish were fully anesthetized (checked by tail pinch test), fish were transferred to a cotton pad soaked with the same concentration of tricaine. A fluorescent dissecting scope was used to visualize the *Nfkb:eGFP* and thin, shallow incisions were made posterior to Nfkb placodes using a razor blade. Fish were imaged using our live imaging setup immediately after incisions were made. Following imaging, fish were revived and placed back on the system.

### Pharmacological Experiments

Fish were imaged on day 1 prior to treatment, then placed in aquarium water with the pharmacological compound or vehicle control diluted to working concentration for the duration of treatment. Fish were maintained off the aquarium system in the dark. For Bay11-7085 and IWR-1-endo, fish were treated continuously in aquarium water off system. Fish were fed and the treatment medium was refreshed once every 24 hours. Bay11-7085 (ApexBio B3033) inhibits IκBα phosphorylation, which prevents NF-κB from entering the nucleus to activate transcription. The efficacy of inhibition was determined by using the *Nfkb:eGFP* reporter line, which showed a loss of eGFP signal after full inhibition. Fish were treated with 0.5μM Bay11-7085 or 0.0005% ethanol in 500ml of aquarium water off system. IWR-1-endo (Cayman Chemical Company 13659) broadly inhibits Wnt signaling through Axin stabilization. Fish were treated with 10μM IWR-1-endo or 0.01% DMSO in 100ml of aquarium water off system.

For BGJ398 (SelleckChem S2183), fish were treated in 100ml of aquarium water off system for one hour per day. Between treatments, fish were fed once every 24 hours and kept in 500ml of water off system. BGJ398 is a pan-FGFR inhibitor. Fish were treated with 1μM BGJ398 or 0.001% DMSO in 100ml of aquarium water off system.

For transient treatments, fish were treated post-imaging on day1 until post-imaging on day 3 (around 48 hours). After washout, fish were rinsed with fresh water and placed back on the system.

### Histology

Whole juvenile fish were fixed using 4% PFA for 48 hours at 4°, then rinsed with PBS and transferred to a 30% glucose solution overnight at 4°. Tissues were frozen in Tissue Freezing Medium (General Data TFM-5) and sectioned by cryostat at 14 μm/section. Slides were rehydrated in PBS with 0.1% Tween and stained with DAPI. Sections were imaged using a 20X oil objective on a Leica Sp8 confocal microscope.

### Image Processing and Quantification

Images from live juveniles were stitched using a custom MATLAB script adapted from De Simone et al^21^. Images of histological sections were assembled using the Stitching plugin in ImageJ^32^. For longitudinal experiments, time points were registered to each other using the imregister function in MATLAB, which aligns images based on image intensity. Registration settings were adjusted as needed for accurate alignment and checked manually. Nfkb or Fgf placodes and scales were counted manually in ImageJ. As images obtained were not always the exact same size, the same size area was used between fish of the same experiment. Scale area and centroid location were obtained manually in ImageJ using the polygon function to outline each scale. Centroid locations were used to construct a Voronoi tessellation in MATLAB. The output of the tessellation was used to determine the number of scales adjacent to a single scale and the distance between neighboring scales.

NF-κB activity was calculated by taking the integral of the GFP intensity profile for *Nfkb:eGFP*. The intensity profile was obtained by measuring the intensity of the *Nfkb:eGFP* channel across a single row of developing placodes along the anterior-posterior axis. When integrating, lines of the same length were used across fish. To determine the distance travelled by the *Nfkb:eGFP* reporter, the posterior edge of the wave or most posterior placode of the bottom row was used to mark the NF-κB wave.

## Acknowledgments

We thank Ken Poss and John Rawls for sharing fish strains. We thank Jim Burris, Lawrence Frauen, and Colin Dolan for zebrafish care. We thank Michel Bagnat, Bernard Mathey-Prevot, David McClay and all members of the Di Talia lab for comments on the manuscript. We thank Priyom Adhyapok, Boris Shraiman and Massimo Vergassola for discussions on the physical mechanisms of skin patterning. We thank Alessandro De Simone for help with computational image analysis. We acknowledge support from the NIH (NIAMS R01-AR076342) and from the Shipley Foundation, Inc. (Program for Innovation in Stem Cell Science).

---

**Figure S1.**
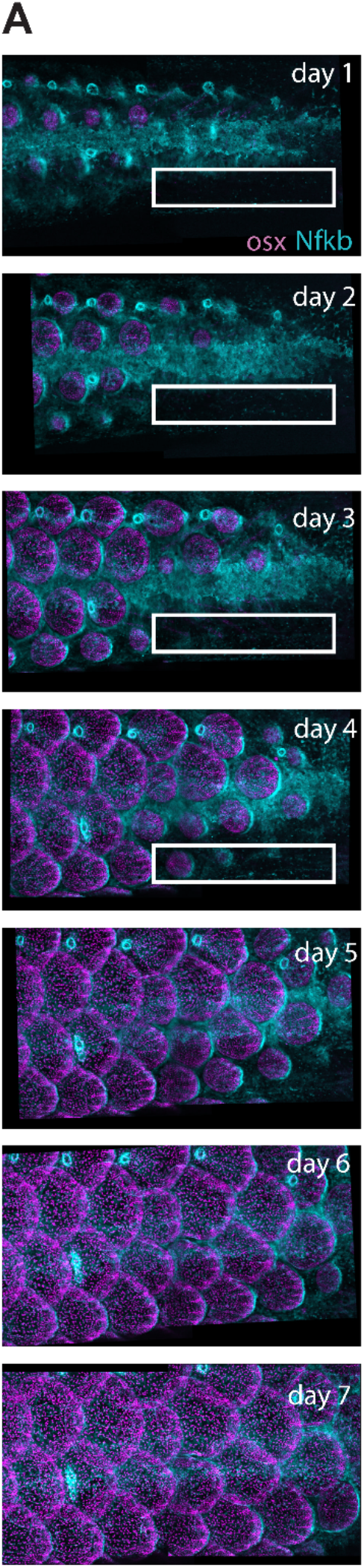
Transverse cuts **A** Representative images of wild-type fish during scale development, cyan: *Nfkb:eGFP*, magenta: *osx-H2A-mcherry*. Small incisions were made in the skin of the fish to block cell-cell communication and test the impact of cut orientation on scale development. The white box outlines the area of the cut. Cuts were made immediately prior to imaging on day 1.

**Figure S2.**
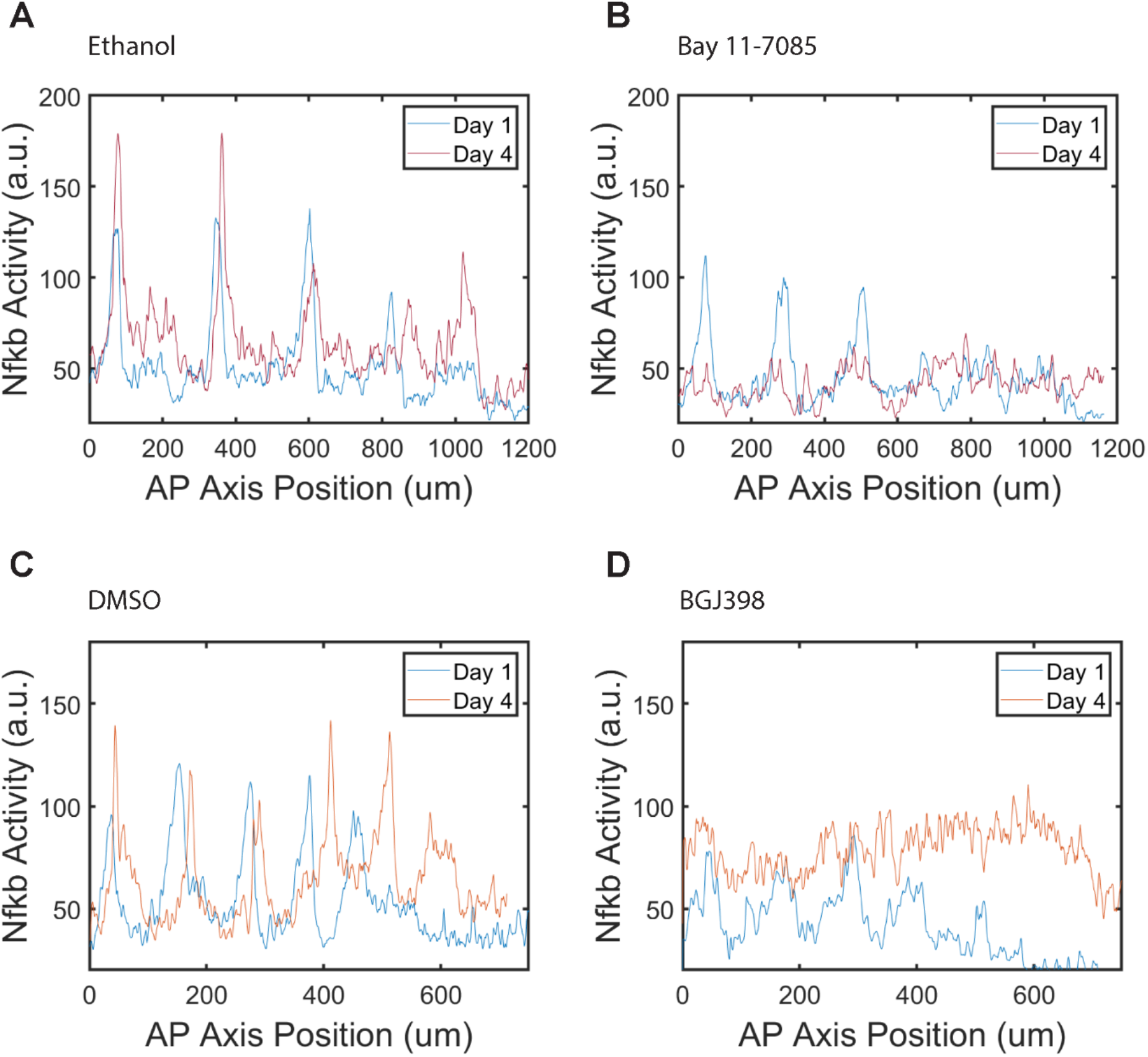
*Nfkb:eGFP* Intensity Profiles **A, B,C,D** The GFP intensity profile was measured across a single row of developing scales parallel to the anterior-posterior axis. The first profile was taken from images before treatment (day 1) and the second profile (day 4) is after approximately 48 hours of treatment.

